# The state of discovery-driven neuroscience research and experimental organism usage in the United States

**DOI:** 10.1101/734798

**Authors:** Sarah M. Farris

**Affiliations:** Department of Biology, West Virginia University

## Abstract

Purely discovery-driven biological research “…performed without thought of practical ends…” establishes fundamental conceptual frameworks for technological and medical breakthroughs that often occur many years later. Despite the critical importance of discovery-driven research for scientific progress, there is increasing concern that it is increasingly less favored by funding agencies than research with explicit goals of application and innovation, resulting in a decline in discovery-driven research output. This in turn appears to promote the use of genetically modified organisms (those with advanced molecular toolkits for gene manipulation and visualization) for which genetic models of human disease can be studied at molecular and cellular resolution using state of the art methodology, and to discourage use of other experimental organisms that provide necessary evolutionary context. This field of neuroscience encompasses both applied and discovery-driven research, providing an opportunity to empirically determine whether funding and publication rates for the latter have indeed declined. Additionally, the diversity of experimental organisms traditionally employed in neuroscience research provides a means to quantify changes in use of study organisms that lack genetic tools over time. In particular, the basic research field of neuroethology is characterized by its distinct approach to selection of study organisms based on their adaptive behaviors, evolutionary history, and suitability for answering the question of interest, providing a stronger basis for the assumption that findings reflect fundamental concepts of nervous system function and behavior. A 30-year analysis of National Science Foundation (NSF) funding of neuroethology research finds that the agency has funded progressively fewer researchers with smaller average award amounts, with a decline in awards for research on non-genetically modified organisms. Neuroscience funding by the National Institutes of Health (NIH) shows the same trend but also increasing support for genetically modified organisms. The same pattern is observed in the neuroscience literature but occurs prior to changes in funding, suggesting that the shift to genetically modified organisms was likely initiated by researchers but may potentially have been later reinforced by funding agency and journal publisher preferences.

## INTRODUCTION

In his historic 1944 report to President Franklin D. Roosevelt titled “Science the Endless Frontier,” Vannevar Bush wrote persuasively of the importance of promoting scientific progress through government funding for basic, discovery-driven research (Bush 1945). Bush’s arguments contributed to a vigorous debate over the role of government in funding scientific research that resulted in the establishment of two federal research funding agencies (the Office of Naval Research and National Science Foundation), and the development of government-funded grant programs for biomedical research through the National Institute of Health (Mazuzan 1994).

Bush defined basic research as that “… performed without thought of practical ends. It results in general knowledge and an understanding of nature and its laws…it provides scientific capital. It creates the fund from which the practical applications of knowledge must be drawn.” The value of basic, discovery-driven research is thoroughly documented by many examples of technological and medical breakthroughs building on fundamental discoveries made years prior, sometimes in seemingly unrelated areas of research (Spector et al 2018; Ronai and Griffiths 2019). As argued by Bush, government support of basic research underlies the United States’ position as a world leader in scientific and technological innovation (National Science Board 2018; Flaherty 2019; Fleming et al 2019).

Despite these facts, there is growing concern that discovery-driven research output in the United States is declining. Decreased federal R&D support, increased emphasis on federally funded STEM research at academic institutions, and growing numbers of scientists entering the academic workforce and applying for federal funding (Couzin and Miller 2007; Rockey 2013; Alberts et al 2014; Harris and Benincasa 2014; Rosbash 2016; Andes and Correa 2017; Mervis 2017) have caused success rates for grants submitted to federal funding agencies to decline (National Science Foundation 2016; NIH RePORT 2019). When the concept of “translational” research was established as a core value of research funded by the NIH (Zerhouni 2003, 2005), it was interpreted as discouraging discovery-driven research that could not be readily applied to clinical outcomes. Although the NIH has maintained that this assertion is not supported by official policy, a perceived benefit to applied, disease-focused research in a highly competitive funding environment has influenced scientists’ choice of research direction (Morrison 2010; Zoghbi 2013; Alberts et al 2014; Kaiser 2014; Landis 2014; Lauer 2016).

A fundamental component of modern biomedical research is the use of non-human animal models (National Institute of General Medical Sciences 2018). The Human Genome Project launched in 1990 with the promise of using the sequenced human genome to improve understanding of the genetic basis of human health and disease, and to provide a foundation for development of molecular targets for diagnosis and treatment (Committee on Mapping and Sequencing Human Genome, National Research Council 1988; Green et al 2011). Gene homology across living organisms provided a rationale for developing animal models of the genetic bases of human health, leading the Human Genome Project to initiate genome sequencing for five model organisms including the roundworm *Caenorhabditis elegans*, the fruit fly *Drosophila melanogaster*, and the mouse *Mus musculus*. These species were already established as experimental organisms in genetics research (Davis 2004) and were (and still are) supported by large communities of scientists that maintain and share resources. Additionally, these communities continuously develop, refine, and distribute cutting-edge technologies for manipulating gene expression and cell function and for visualizing processes at the cellular and subcellular level. Today, these genetically modified organisms are powerful tools for mechanistic studies of gene function in fundamental biological processes and for developing new gene-based technologies that could enhance understanding of human health and treatment of disease. As such, they play a prominent role in biomedical research supported by the National Institutes of Health (National Institute of General Medical Sciences 2018).

In 1999, the NIH released a publication containing a “canonical list” of model organisms (the original 1999 online publication “Non-mammalian models workshop” is no longer available online, but Dietrich et al (2014) provides the list of organisms). Animals on the canonical list included the three “genetic model organisms” listed above in addition to a newly developed model, the zebrafish *Danio rerio* (“zebrafish”). Three model organisms without advanced genetic toolkits were also included on the list: the African clawed frog species *Xenopus laevis* or *X. tropicalis* (“*Xenopus*”), the chicken *Gallus gallus* (“chick”), and the rat *Rattus rattus* (“rat”). This announcement was met with skepticism as scientists became concerned with the increasingly narrow focus on molecular mechanisms in a handful of species that this announcement seemed to promote. For example, while genetic model organisms provide unparalleled ability to experimentally manipulate gene and cell function, they all have unique features that are not entirely representative of fundamental biological processes. Caveats to the use of these organisms include the fact that most were selected for unusual traits such as rapid and canalized development (Bolker 1995) and have been laboratory bred for generations so as to reduce genetic variation and exposure to environmental factors that occur in nature (Brenowitz and Zakon 2015; Bolker 2017). There is also increasing concern that the focus on technological advances drives an excessively reductionistic approach to research such that the relationship between the narrowly defined, cellular, and behavioral preparation and their function in the whole organism is no longer clear (Bolker 2017; Krakauer et al 2017). As stated by Bolker (2012), “The extraordinary resolving power of core models comes with the same trade-off as a high-magnification lens: a much reduced field of view.” In order to judge the results of mechanistic studies as representative of fundamental biological processes, they must be considered in the context of their adaptive function in the whole organism. This context has traditionally been provided by discovery-driven research in non-genetic experimental organisms that provide exceptional examples of a given biological process but lack genetic tools. There are many instances of the integration of these two approaches providing important insight into processes with potential impact on human health; for example, the role of the mammalian cerebellum in predicting sensory consequences of movement benefited from years of detailed neurophysiological studies of electrosensory processing by a cerebellum-like structure in weakly electric fish (reviewed by Nixon 2003; Warren and Sawtell 2016; Sawtell 2017). Studies in a wide variety of species can also demonstrate that discovery made in a genetic model species is unique to that species, rather than being representative of a fundamentally conserved biological process (Brenowitz and Zakon 2015).

Is purely discovery-driven neurobiological and behavioral research, as exemplified by neuroethology on the decline? If so, is the decline associated with increased usage of genetic model species? The field of neuroscience provides an opportunity to answer this question as it traditionally includes a wide variety of disciplines ranging from purely discovery-driven to applied, clinical research. One classically discovery-driven field is that of neuroethology, which has a long and rich history with numerous contributions to neuroscience including Nobel Prize-winning discoveries. Neuroethology is defined as the study of the neural basis of behavior, and theoretical and methodological aspects of animal behavior (ethology), comparative neuroanatomy, and neurophysiology (Ingle and Crews 1985; Pfluger and Menzel 1999). Of particular relevance to this study, neuroethological research is characterized by the use of experimental organisms whose natural behaviors are particularly well adapted to solve the problem of interest to the researcher (Camhi, 1984; Hoyle 1984). As such, species of interest are selected based on their natural behaviors, evolutionary history, and experimental tractability, resulting in the use of diverse species spanning the animal kingdom from primates to jellyfish. This approach is fundamentally different from that of more applied biomedical research in which the experimental organism may be selected for the availability of genetic tools and the ability to engineer the models of human health and disease (Bolker 2017).

In this study, funding, publications, and experimental organism usage in neurosciences are tracked over the past 20-30 years, depending on data availability. Previous studies of the association between funding, publications, and model systems usage have focused on NIH funded research which supports both discovery-driven and disease-focused studies. This study’s primary focus is on funding by National Science Foundation (NSF) which as described in The National Science Foundation Act of 1950 “…is authorized and directed…to initiate and support basic scientific research” (81^st^ United States Congress, 1950). The mission of the National Science Foundation is thus more explicitly focused on funding discovery-driven research than the NIH, although it has a far smaller budget (6.487 billion US$ vs 36.097 billion US$ in 2018; AAAS 2019). The Integrative Organismal Sciences (IOS) division of the NSF has traditionally supported neuroethology research employing an array of experimental organisms through several funding programs (see Methods). However, this support has declined precipitously as proposal submissions have increased, as has also been the case for NIH funding (Rockey 2013). neuroethology investigators are increasingly funded through smaller collaborative awards, and grants for non-genetic model organism research have decreased. All measures of funded awards and publications reveal that research on experimental organisms lacking genetic toolkits has decreased, while that for genetically modified organisms has remained steady or increased. However, the proportions of genetic vs non-genetic experimental organisms began to change in the publication record prior to changes in funding, supporting previous observations that the research community itself initiated the rise to dominance of genetically modified organisms (Landis 2014; Pierson et al 2017).

## METHODS

### Neuroscience and behavioral research funding

The NSF Award Search advanced search engine (https://www.nsf.gov/awardsearch/advancedSearch.jsp) was used to gather award data from the BIO Directorate, the Division of Integrative Biology and Neuroscience (IBN; 1987-2004), Division of Integrative and Organismal Biology (IOB; 2004-2007), and the Division of Integrative Organismal Systems (IOS; 2007-present) every other fiscal year (Oct 1-Sept 30) from 1987-2017. Clusters that fund neurobiological and behavioral research (considered together as neuroethology) were filtered from the above divisions. From FY 1987-2004, the Neuroscience cluster was composed of the Behavioral Neuroscience, Computational Neuroscience, Developmental Neuroscience, Neuroendocrinology, Neuronal and Glial Mechanisms, and Sensory Systems programs. The Animal Behavior program was contained within the Physiology and Ethology cluster. From 2004-2007, neuroscience-focused programs at NSF were reorganized into the Environmental & Structural Systems and Functional and Regulatory Systems clusters. The Animal Behavior program was placed within the Behavioral Systems Cluster where it has remained through the present. In 2007, the Neural Systems cluster was established and subdivided into the Organization, Activation, and Modulation programs which remain today. It should be noted that the transition between IBN, IOB, and IOS clusters overlapped; for example, an award granted in 2007 might be made through both the older Environmental and Structural Systems Cluster and the Activation program (part of the newer Neural Systems Cluster). Documentation of the organization of these divisions and clusters was obtained from the NSF Document Library (https://www.nsf.gov/publications/).

NSF BIO, IOS, and neuroethology award amount data was converted to 2018 United States dollars ($) using the CPI Inflation Calculator provided by the US Bureau of Labor Statistics (https://data.bls.gov/cgi-bin/cpicalc.pl). Dollar amounts were analyzed only through 2011 as the available data provided only amounts awarded to date, resulting in incomplete totals for ongoing awards. Total numbers of grants awarded and PIs funded were analyzed up to 2017. Total numbers of proposals submitted to IOS and grants awarded from 2001-2018 were collected from https://dellweb.bfa.nsf.gov/ and https://catalog.data.gov/dataset/nsf-research-grant-funding-rate-history. Numbers of preliminary proposals submitted and accepted to IOS were obtained from Katz et al (2017).

Model organisms used in funded NSF neuroethology research was determined from a curated search of abstracts for each funded neuroethology award and, if necessary, from a PubMed search (https://www.ncbi.nlm.nih.gov/pubmed) of publications by the PI at the time of the award. Genetically modified model organisms were defined as animals for which tools are available for germ-line transformation and modification of gene expression that were also named as “canonical” model organisms by the National Institutes of Health in a 1999 online publication titled “Non-mammalian models workshop.” This document is no longer available online, but Dietrich et al (2014) provides the list of organisms. Organisms considered genetic models were the roundworm *Caenorhabditis elegans*, the fruit fly *Drosophila melanogaster*, the mouse *Mus musculus*, and the zebrafish *Danio rerio*. All other animal species used in funded research, as well as computational research, were grouped as “non-genetic model” organisms.

‘The NIH RePORT search engine (https://report.nih.gov/index.aspx) was used to collect data on research project grants (RPGs) awarded by NIH neuroscience programs. Institutes participating in the NIH BRAIN Initiative were included in the search (Koroshetz et al, 2018). Awards were searched for use of triploblastic animal model organisms on the canonical organisms list. Genetic model organisms are those described above while non-genetic canonical model organisms were the chicken *Gallus gallus*, the African clawed frog *Xenopus laevis* or *X. tropicalis*, and the rat *Rattus rattus*). The search for award descriptions containing these organisms was performed for every other year from 1995-2017 using the following search strings: *Drosophila* or *melanogaster* or “fruit fly” or fruitfly; zebrafish or *Danio* or *rerio*; *Caenorhabditis* or *elegans*; rat or *Rattus*; mouse or *Mus* or *musculus*; chick or chicken or *Gallus*; (frog or *Xenopus* or *laevis* or *tropicalis*) NOT oocyte NOT egg.

### Neuroscience and behavior research publications

A Web of Science advanced search (https://apps.webofknowledge.com) was used to identify journal articles published in the Neurosciences subject area by year, model species, and United States authorship. Publication searches comparing NIH-designated genetic and non-genetic model organisms were restricted to article titles. This method underreports numbers of publications with the term of interest (Dietrich et al 2014), but avoids false positives caused by abstract and keyword mentions of species not used in the study. Curated Web of Science topic searches were performed for comparisons of two sets of genetic model and non-model organisms: the fruit fly *Drosophila melanogaster* vs. insects (all other insect species) and the zebrafish *Danio rerio* vs. fish (all other fish species).

Data was plotted using Microsoft® Excel® for Mac 2011 (Microsoft Corporation, Redmond WA, USA) and fit of regression lines was determined via ANOVA using the GraphPad QuickCalcs browser application (GraphPad Software, San Diego CA, USA) (https://www.graphpad.com/quickcalcs/).

## RESULTS

### Neuroethology research funding by the National Science Foundation

Animal behavior and neuroscience research (here considered together as neuroethology) is funded by the NSF under the Division of Integrative and Organismal Systems (IOS; previously designated the Division of Integrative Biology and Neuroscience or the Division of Integrative and Organismal Biology) that is in turn contained within the BIO directorate. Funding through these bodies was determined via an advanced award search for every other fiscal year from 1987-2011, after which complete funding data was unavailable due to ongoing awards. When converted to 2018 US dollars, BIO funds (Figure 1A; F(1, 11)= 35.78, p<0.0001, R^2^= 0.7648) and IOS funds(Figure 1A; F(1,11)= 19.05, p=0.0011, R^2^= 0.6340) disbursed has increased significantly. From 1987-2017 the total number of awards made by both the BIO directorate (Figure 1B; F(1, 14)= 0.4856, p=0.4973, R^2^= 0.0335) and the IOS division (Figure 1B; F(1, 14)= 1.113, p=0.3094, R^2^= 0.0736) remained constant. Thus, on average, the dollar amount per award increased from 1987-2011 for grants made through the BIO directorate and the IOS division.

**Figure 1.**
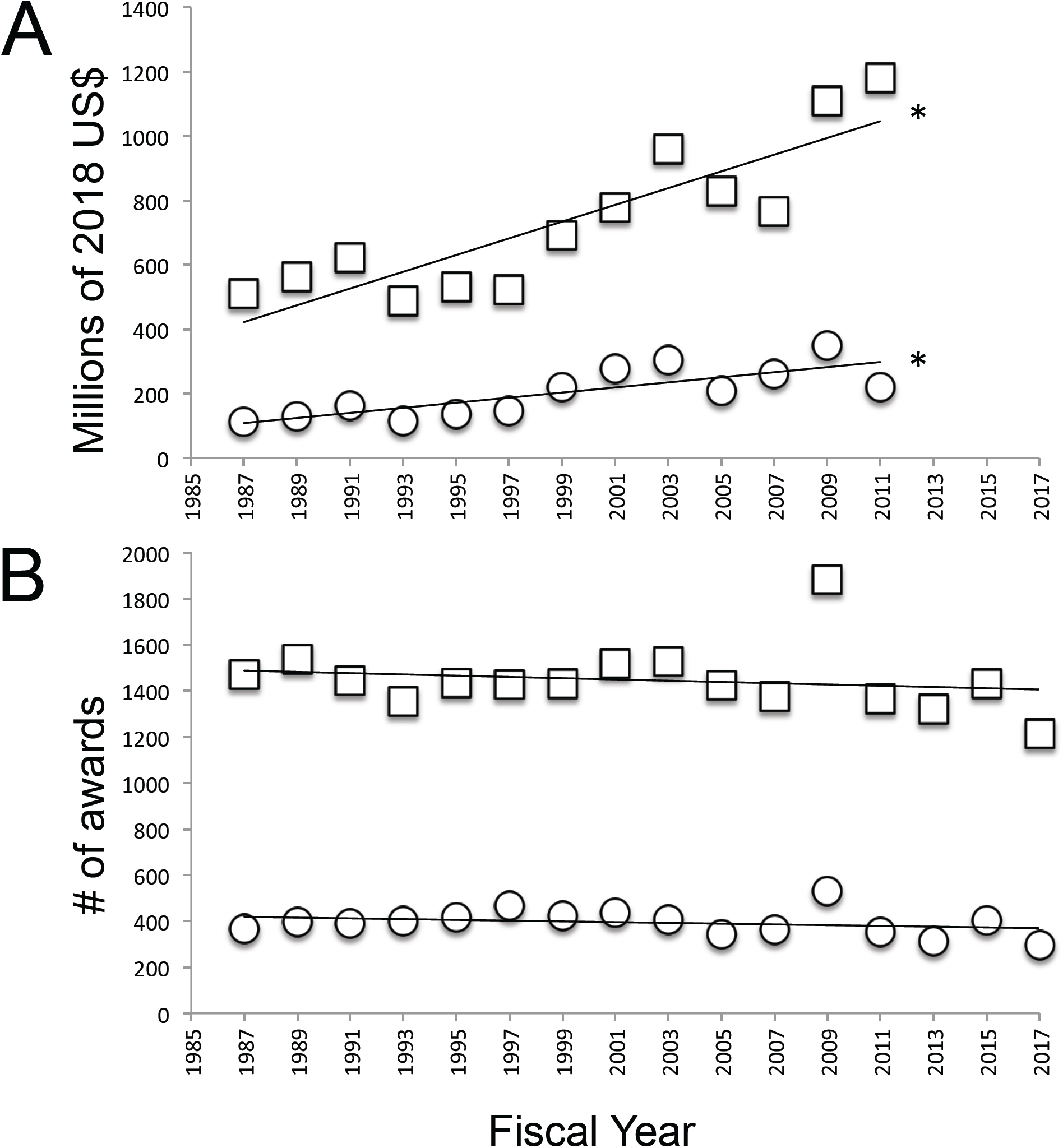
A. Total 2018 US$ awarded every other year from FY1987-FY2011 by the BIO directorate (squares) and IOS division (circles). B. Total award numbers made every other year from FY1987-FY2017 by the BIO directorate (squares) and IOS division (circles). Asterisks indicate significant change over time.

Beginning in 1998, grants funded through the Plant Genome Project (PGP) were added to the IOS Division (MacIlwain 1997). When these grants are subtracted from all grants funded through IOS (IOS-PGP), the amount of funding disbursed via IOS did not change from 1987-2011 (Figure 2A; F(1, 11)= 2.635, p=0.1328, R^2^= 0.1933), as was also the case for neuroethology awards funded through IOS programs (Figure 2A; F(1, 11)= 0.1099, p=0.7465, R^2^= 0.0099). The number of IOS-PGP awards decreased from 1987-2017 (Figure 2B; F(1, 14)= 5.736, p=0.0312, R^2^= 0.2906), again mirrored by a decrease in NBR awards made (Figure 2B; F(1, 14)= 28.29, p=0.0001, R^2^= 0.6690). As observed for BIO and IOS, IOS-PGP and neuroethology awards were fewer and larger; however, in the case of IOS-PGP and neuroethology this occurred due to a decline in the number of awards made and a flat budget.

**Figure 2.**
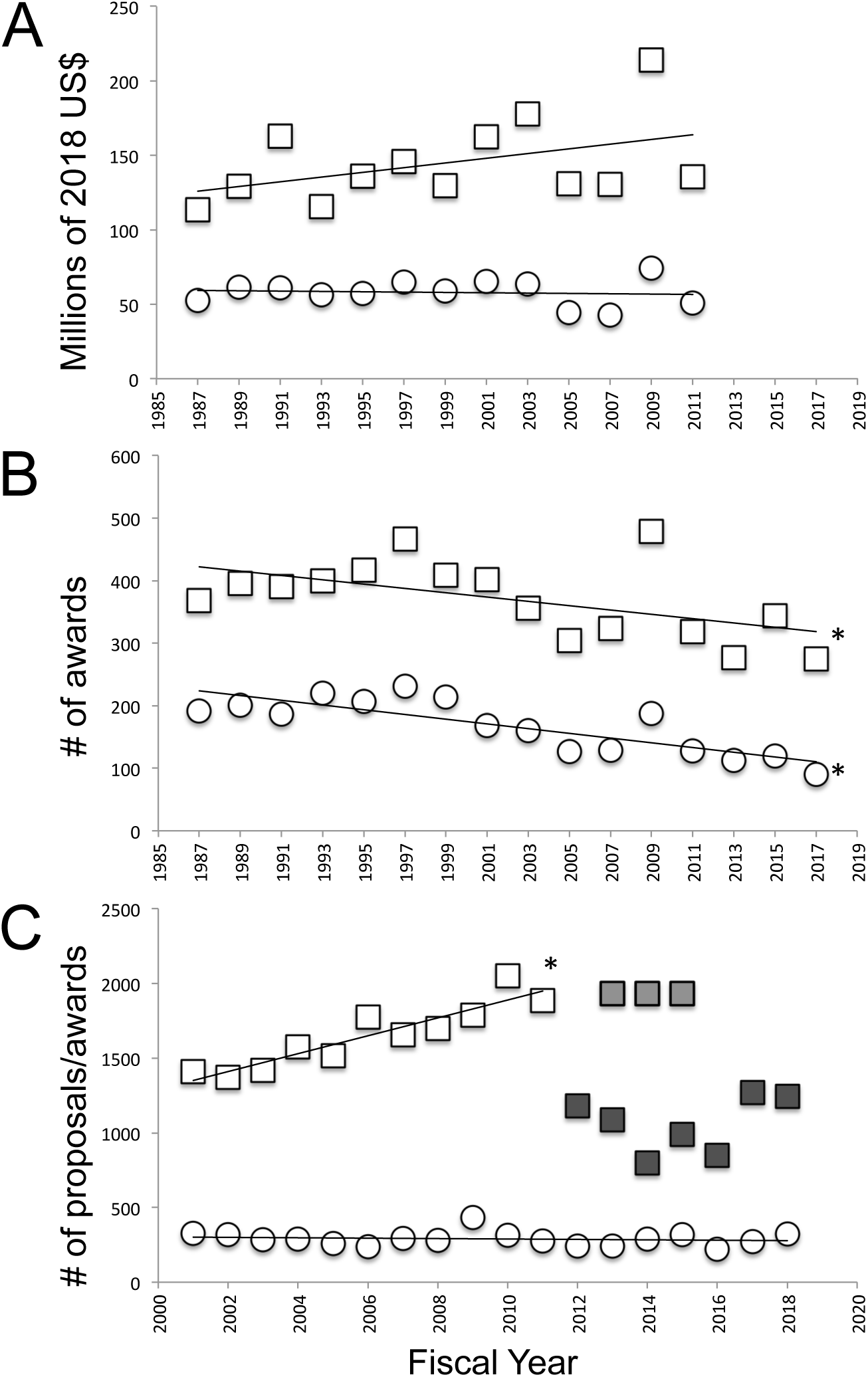
A. Total 2018 US$ awarded every other year from FY1987-FY2011 by the IOS division (Plant Genome Project awards subtracted (IOS-PGP); squares), and neuroethology programs contained within IOS (circles). B. Total award numbers made every other year from FY1987-FY2017 by IOS-PGP (squares) and neuroethology programs (circles). C. Full proposals submitted to IOS from 2001-2011 (open squares), estimated preproposals submitted from 2013-2015 (light grey squares), invited full proposals submitted from 2013-2015 (dark grey squares) and total grants funded from 2000-2018 (circles). Asterisks indicate significant change in award number over time.

Do decreasing numbers of awards funded reflect a decrease in proposals submitted? Submission data specifically for neuroethology or IOS-PGP was not available; however, data for IOS from 2001-2018 revealed a steady increase in proposal submissions from 2001-2011 (Figure 2C; F(1, 9)= 49.87, p<0.0001, R^2^= 0.8471), while the number of grants awarded remained unchanged (F(1, 16)= 0.3850, p=0.5437, R^2^= 0.0235), resulting in a steadily declining success rate. In 2012 IOS adopted a preliminary proposal requirement (Roskoski 2011). These short proposals were evaluated and a portion of the submitting PIs chosen to submit full proposals. Katz et al (2017) reports that from 2013-2015 a total of 5802 preproposals were submitted to IOS; divided equally across the three-year period, preproposals submitted per year from 2013-2015 were likely to be roughly the same as the number of proposals submitted prior to 2012 (grey squares, Figure 2C). Katz et al (2017) further report that of 5802 preproposals submitted to IOS, 1344 were chosen for submission as full proposals. While the number of awards made out of invited full proposals appears to be a dramatic improvement over success rates of previous years (Figure 2C, black squares), considering awards made out of the number of preproposals reveals a similar success rate to that prior to implementation of the preproposal requirement. Although this data contains all of the programs that are part of IOS and not just those funding neuroethology, it suggests that the observed decrease in neuroethology grants funded may not necessarily result from a decrease in submitted proposals.

The decrease in total neuroethology awards particularly impacted the number of sole PI research awards (excluding conference, travel, and dissertation grants) (Figure 3A; F(1, 14)= 98.01, p<0.0001, R^2^= 0.8750). From a peak of 160 sole PI awards in 1993, only 30 sole PI awards were granted in 2017, a decline of over 80%. In contrast, little change in the number of collaborative awards made from 1987-2017 was observed (Figure 3A; F(1, 14)= 1.120, p=0.3078, R^2^= 0.0741). As sole PI awards have decreased, increasing numbers of researchers were funded on collaborative grants (Figure 3B). On average, PIs on collaborative grants received smaller awards than did sole PIs, even though mean award amounts increased for all PIs from 1987-2011 (Figure 3C). To summarize, successfully funded NSF neuroethology grants have declined both in number and in average amount awarded.

**Figure 3.**
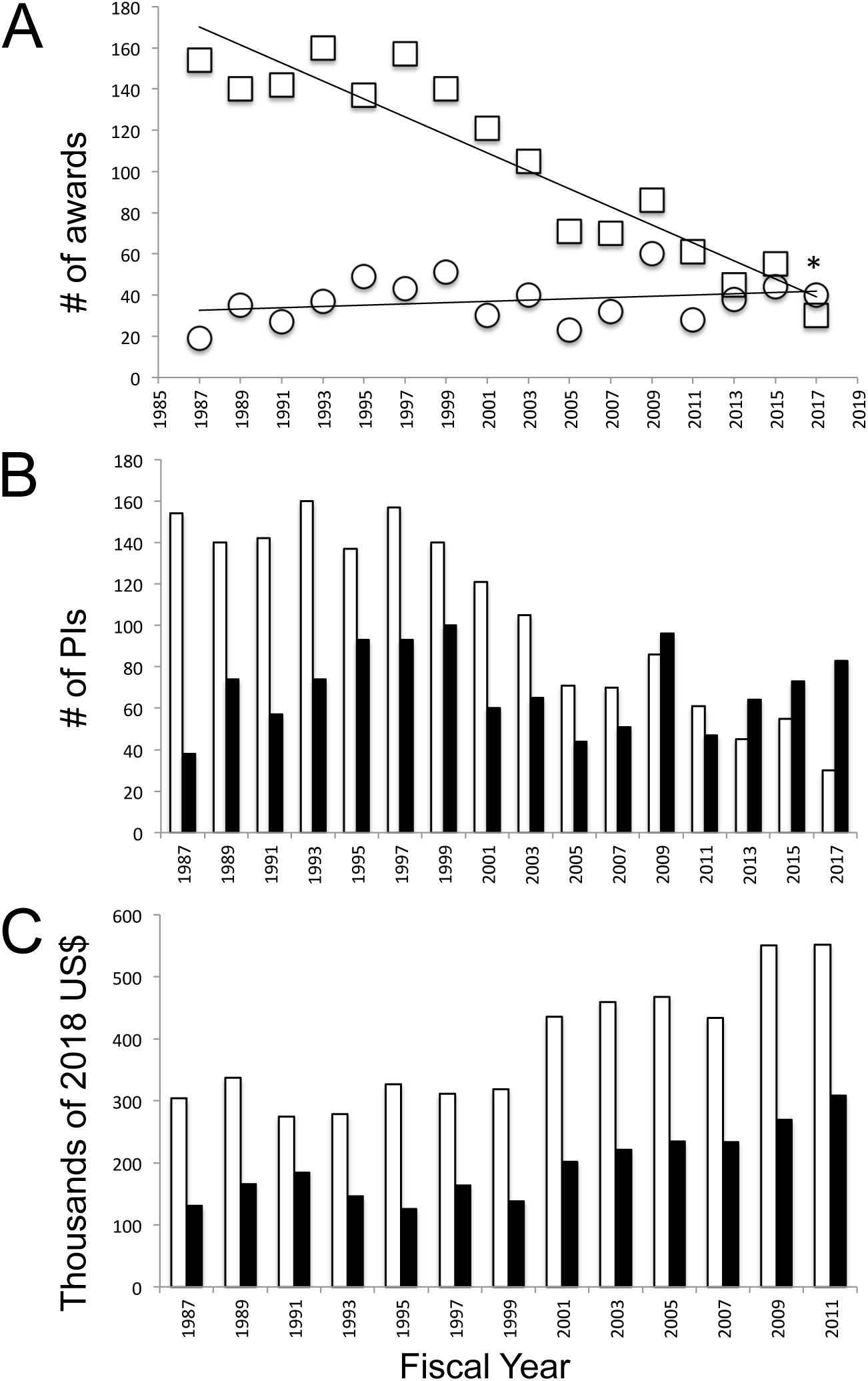
A. Sole PI (squares) and collaborative (circles) neuroethology research awards from FY1987-FY2017. B. Numbers of individual sole PIs (white) and individual PIs funded on collaborative grants (black). C. Mean award amounts for sole PIs (white) and PIs on collaborative grants (black bars). Asterisk in A indicatse significant change over time.

### Model organism usage in neuroethology research awards and publications

From 1997-2017, the decline in total NSF neuroethology grants funded particularly impacted the number of awards proposing research using non-genetic model organisms (Figure 4A; F(1, 9)= 49.13, p<0.0001, R^2^= 0.8452). The non-genetic model organisms included in this measure were those designated as canonical models by the NIH but that lack molecular tools (chick, African clawed frog, rat) as well as many other species traditionally used in neuroethological research such as electric fish, songbirds, and honey bees. NSF neuroethology awards for genetic model organism research remained low but constant over this time period (Figure 4A; F(1, 9)= 3.286, p=0.1033, R^2^= 0.2675). Due to the decline in awards for non-genetic model organisms, however, those for genetic model organisms made up almost 28% of total NSF neuroethology research awards in 2017, compared with 17% in 1997.

**Figure 4.**
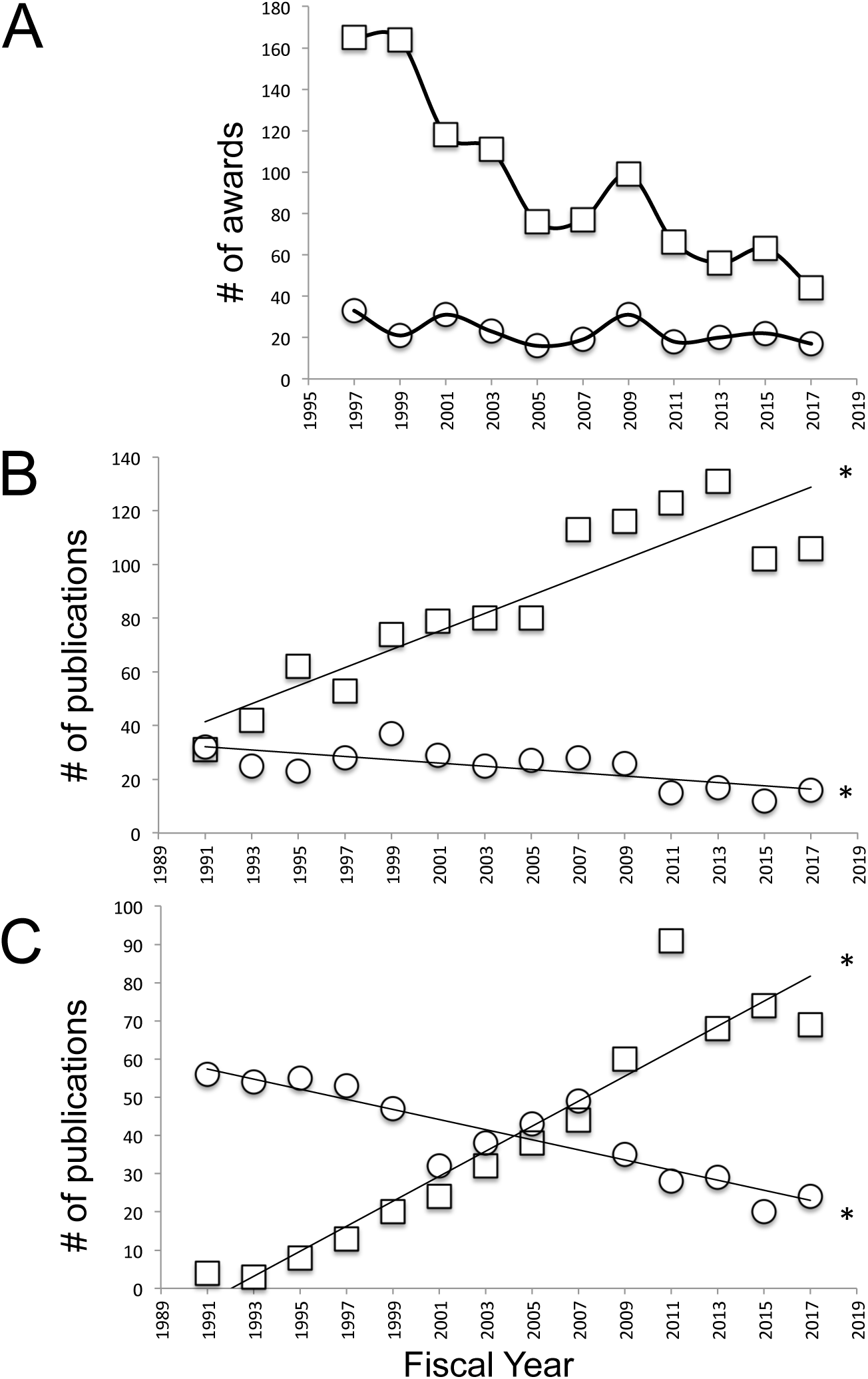
A. NSF neuroethology awards for non-genetic model species research (squares) and genetic model species research (circles) from 1997-2017. B. Curated publications by United States authors using the Drosophila model species (squares), US authors using other insect model species (circles), and authors from other countries using other insect model species (triangles). C. Curated publications by United States authors using the zebrafish model species (squares), US authors using other insect model species (circles), and authors from other countries using other insect model species (triangles). Asterisks indicate significant change in publication number over time.

Changes in funding of non-genetic model organism (by both the NSF and NIH, as shown below), mirrored changes in neuroethology publications. Two curated Web of Science topic searches of US authored neuroethology publications compared large phylogenetic groups of organisms (insects and fish; Figures 4B and 4C respectively) that also contained a genetic model organism (*Drosophila melanogaster* and the zebrafish, respectively). Research articles on any non-genetic model species of fish or insect have significantly declined since 1991, while publications employing the genetic model species have significantly increased (Figure 4B, *Drosophila*; F(1, 12)= 54.00, p<0.0001, R^2^= 0.8182: Figure 4B, all other insects; F(1,12)= 13.04, p=0.0036, R^2^= 0.5209: Figure 4C, zebrafish; F(1, 12)= 98.76, p<0.0001, R^2^= 0.8917: Figure 4C, all other fish; F(1, 12)= 46.77, p<0.0001, R^2^= 0.7958).

NIH RePORTER was used to search for awards granted for research on each animal canonical model species by neuroscience-focused institutes. Awards for genetic model organism research increased and that for non-genetic model organisms decreased after 1999 (Figure 5A). Web of Science title searches by model organism revealed opposing trends for the two rodent model species: steadily increasing numbers of publications for the genetic model (mouse) and declining publications for the non-genetic model (rat), similar to that reported by Dietrich et al (2014; data not shown). Web of Science publication searches revealed increasing numbers of publications for the remaining animal genetic model species (*C. elegans, Drosophila melanogaster*, and zebrafish) and decreasing numbers of publications for the non-genetic model species (*Xenopus* and chick). Unlike the trends in NIH award funding after announcement of the canonical model species in 1999 (Figure 5A), changes in publication rates began prior to 1999.

**Figure 5.**
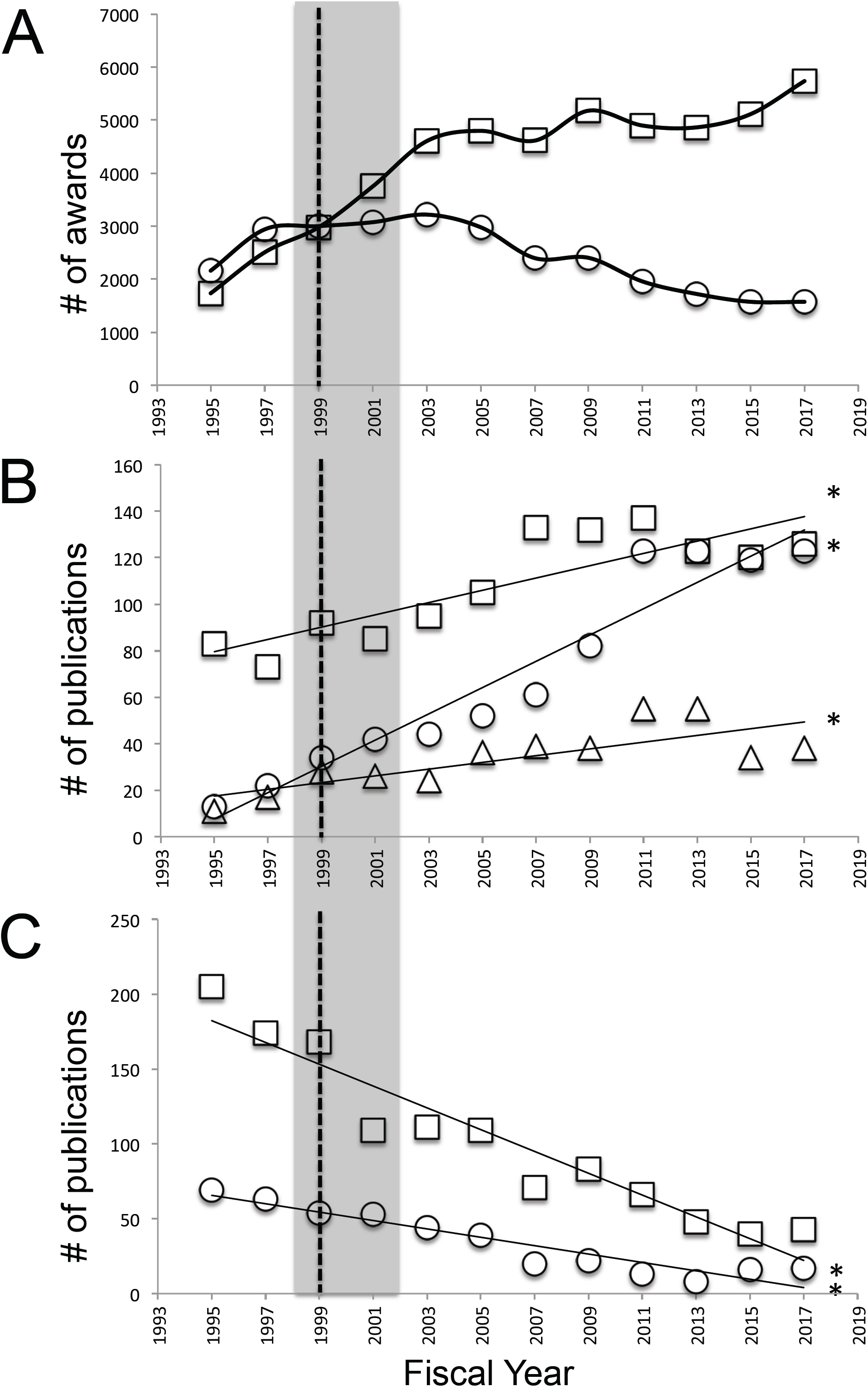
A. NIH neuroscience awards for canonical animal model species with (squares) and without (circles) genetic tools. B. Title search for publications using canonical genetic model species (mouse, zebrafish, *Drosophila, C. elegans*). C. Title search for publications using canonical non-genetic model species (*Xenopus*, chick). Dotted line indicates release of NIH list of canonical model species in 1999. Shaded area indicates time period in which genome sequences were completed for three genetic model species, the mouse, *Drosophila*, and *C. elegans*.

## DISCUSSION

This study sought to answer two questions about the state of research in the field of neuroscience. First, is purely discovery-driven neurobiological and behavioral (neuroethological) research in the United States on the decline? The neuroscience subfield of neuroethology is defined by used of a comparative approach, selecting species for their adaptive behaviors that address the question of interest, allowing the neural mechanisms of these behaviors to be studied. Two measures were used to determine the trajectory of this field of research: funding by the National Science Foundation (NSF; 81^st^ United States Congress 1950), charged with supporting discovery-driven research without explicit applications, and experimental organism usage in published neuroscience literature. Here, a decline in funding for awards granted by NSF neuroethology programs and a decline in non-genetic model organisms/increase in genetic model organisms in the general neuroscience literature is interpreted as a decline in the traditional neuroethological approach to neuroscience research.

The second goal was to determine whether a small number of experimental organisms for which advanced molecular tools are available (genetic model organisms) have come to dominate funded and published neuroscience research as a whole. This question reflects concerns that the National Institutes of Health (NIH) has increasingly emphasized “translational” or “bench to bedside” research that is explicitly disease-focused (Zerhouni 2003; Morrison 2010; Zoghbi 2013; Alberts et al 2014), and as such increasingly focused on a small number of NIH designated “canonical” model organisms for which highly conserved genetic pathways can be studied using state of the art molecular tools available in genetic model organisms. It is important to state that purely discovery-driven research also benefits from the use of genetic model organisms, however, generalization of deeply reductionist studies in a handful of species must be balanced the evolutionary and adaptive context provided by organismal and species-level studies such as those characteristic of neuroethology (Bolker 2017; Krakauer et al 2017).

The analysis of NSF award data from 1987-2017 demonstrates a decline in support by this agency for discovery-driven neuroethological research. While total funds awarded by the BIO directorate and IOS division have increased and the number of awards has remained stable (Figure 1), funds awarded by neuroethology funding programs contained within IOS were stagnant, and the number of awards made decreased (Figure 2A, 2B). The magnitude of the decline in grants and researchers awarded is striking: in 2017, only 70 research awards and 113 PIs were awarded funding for discovery-driven neuroethology research, 65% and 55% of that in 1987, respectively (Figures 2B, 3A). Although data on the number of proposals submitted to NSF neuroethology programs was not available, total submissions to IOS increased from 2001-2011 (Figure 3C). A similar trend occurred at the NIH, where increasing numbers of grant submissions and constant numbers of awards made resulted in decreasing success rates (Rockey 2013; NIH RePORT 2019). It is thus reasonable to infer from this data that proposal submissions to IOS neuroethology programs also increased.

The NSF neuroethology data differs from that for IOS and NIH, however, in that the number of awards made significantly decreased from 1987-2017 (Figure 2B) rather than remaining constant. NSF neuroethology programs also tended to fund an increasing proportion of investigators on collaborative grants, for which award amount to each PI was consistently lower than that for sole PI grants (Figure 3). Thus, regardless of changes in the numbers of proposals submitted to NSF neuroethology programs, there has been a trend towards funding fewer proposals and fewer investigators for smaller amounts. Interestingly, the decline in neuroethology awards funded was associated with a decrease in awards for non-genetic model organism research after 1999 (Figure 4A; see below).

The hyperdiverse insects and fish are established neuroethology models due to their rich evolutionary history and natural behavioral repertoire, and the relative ease with which they can be laboratory reared and used in behavioral, anatomical, and neurophysiological research. Each of these large taxa includes a genetic model organism: the fruit fly *Drosophila melanogaster* and the zebrafish, respectively. Curated searches of United States neuroscience publications demonstrate that genetic model species usage has increased while usage of all of the non-genetic model species in that taxon has decreased (Figure 4B, 4C). Together, these data suggest that the field of neuroscience, including neuroethology, is increasingly focused on a few genetic model species.

In 1999 the NIH released a publication designating “canonical” model organisms, causing concern that research on organisms not included on the list would be discouraged (Dietrich et al 2014). Analysis of grants funded by NIH neuroscience institutes (determined by their participation in the BRAIN Initiative) shows that funded awards for canonical model organisms did increase after 1999, but only for those species with genetic tools. Canonical model organisms lacking these tools such as the rat, chick, and frog, were used in progressively fewer funded grants (Figure 5A). Interestingly, the decline in NSF-funded awards for non-genetic model species also occurred after 1999 (Figure 4A), although the canonical model species designation did not apply to the NSF. Increasing usage of genetically modified canonical organisms and decreasing usage on non-genetic organisms was also observed in the neuroscience literature but predated the 1999 increase genetic model funding (Figure 4B, 4C, 5B, 5C). This interpretation supports NIH data demonstrating that grants are not preferentially awarded for canonical and/or genetic model species research (Willis and Basson 2018; Lauer 2016a; Lauer 2016b), although this data does not include the years immediately before and after the designation of the canonical model species.

Taken together, the data presented here suggests that there has indeed been a shift in model organism usage in neuroscience, away from non-genetic organisms and towards genetically modified organisms. This shift began in the literature, suggesting that it was initiated by researchers or by a preference for genetic model studies and perhaps their perceived superior impact by journal editors and reviewers (for example see Steinberg et al. 2016) and eventually manifested in changes in model organisms used in funded research. Bolker (2017) suggests that a self-perpetuating cycle is maintained by the communities of researchers that form around these genetic model species, as they become convinced of the superiority of that organism and subsequently influence funding, publication, and training of the next generation of scientists. Thus a combination of factors may have propelled genetic model organisms to their dominance in neuroscience research.

What caused the initial shift to genetic model organisms occur, if not explicit grant funding incentives? As described above, the predominant genetic model species are supported by communities of researchers, facilitating the development and transmission of new technologies (Davis 2004). The rise in neuroscience publications using genetic model organisms began in the 1990s, a period of rapid refinement and adoption of key molecular tools such as targeted recombination, germ line transformation, and spatial and temporal control of gene expression for *Drosophila* and mouse, culminating in the release of genome sequences for these and other genetic model species (Gordon and Ruddle 1981; Rubin and Spradling 1982; Thomas and Capecchi 1987; Golic and Lindquist 1989; Orban et al 1992; Brand and Perrimon 1993; International Human Genome Sequencing Consortium, 2004). Publication of these advances in top tier journals and the promise of new approaches and areas of study enabled by these molecular tools likely attracted many researchers to these model system communities, thus increasing representation of these organisms in the literature. However, it is possible that the continuing increase in usage of genetic model species and decrease in non-genetic model species was influenced by declining grant success rates in the early 2000s. Declining grant success rates at the NSF may have driven neuroethology researchers to submit proposals to the NIH. However, declining success rates at the NIH and emphasis on “translational” research during this time may have further accelerated the shift to genetic model species. These organisms were likely perceived as better able to attract funding due to the ability to invoke gene homology to draw parallels between animal models and human health and disease, and to utilize “innovative” approaches employing state-of-the-art genetic tool that are often specifically indicated in funding announcements. These suppositions might be tested in a future study comparing funding history and model organism usage of individual PIs in the NSF grant database and the NIH RePORTER database.

Hypercompetition for grant funding resulting from stagnant biomedical research budgets and growing numbers of funding applicants is increasingly acknowledged as a major problem facing biomedical scientists in the United States. Most alarmingly, the funding crisis ends careers, promotes making disingenuous claims about the proposed research to meet funding criteria, and requires researchers to spend significant time crafting competitive grant proposals as opposed to performing actual scientific research (Couzin and Miller 2007; Morrison 2010; Alberts et al 2014; Harris and Benincasa 2014; Rosbash 2016). Additionally, the need to maximize funding potential favors depth over breadth: emphasis on “translational,” “transformative,” and “innovative” approaches are interpreted (correctly or not) to favor proposals for more applied research using a handful of experimental organisms for which state of the art genetic toolkits are available, and to which research questions are tailored to the capabilities of the organism such that the model may, at times, poorly represent the actual process being studied (Bolker 2017; Reardon 2018). In contrast, purely discovery-driven research disciplines as exemplified by the field of neuroethology takes a different approach, in which experimental organisms are chosen for adaptive natural behaviors in the area of interest, and are considered within a comparative evolutionary context (Camhi, 1984; Hoyle 1984). Combining this approach with the ability to precisely manipulate genetic and cellular components in a genetic model organism provides powerful insight into complex behaviors in context, from the molecular to the organismal level, and allows novel, convergent, or conserved behavioral mechanisms to be identified (Brenowitz and Zakon, 2015; Bolker 2017). The importance of this dual approach has become increasingly evident as drugs fail after succeeding in mouse models that poorly represent human physiology or disease processes (Perrin 2014; Makin 2018; Reardon 2018).

It is clear that a single genetic model organism cannot be treated as a perfect representative of animal biological processes and it is critical that funding agencies remain cognizant of the importance of discovery-driven research incorporating diverse species. However, in the current hypercompetitive funding climate federal funding agencies must not only support but also incentivize scientists to use non-genetic model organisms that may currently be perceived as less likely to attract funding that is essential for academic career success.

